# Comparison of GWAS models to identify non-additive genetic control of flowering time in sunflower hybrids

**DOI:** 10.1101/188235

**Authors:** Fanny Bonnafous, Ghislain Fievet, Nicolas Blanchet, Marie-Claude Boniface, Sébastien Carrère, Jérôme Gouzy, Ludovic Legrand, Gwenola Marage, Emmanuelle Bret-Mestries, Stéphane Munos, Nicolas Pouilly, Patrick Vincourt, Nicolas Langlade, Brigitte Mangin

**Affiliations:** LIPM, Universit de Toulouse, INRA, CNRS, Castanet-Tolosan, France; LIPM, Université de Toulouse, INRA, CNRS, Castanet-Tolosan, France; TERRES INOVIA, AGIR, Castanet-Tolosan, France

**Keywords:** Genome-wide association study, sunflower, multi-locus, non-additive effect

## Abstract

Genome-wide association studies are a powerful and widely used tool to decipher the genetic control of complex traits. One of the main challenges for hybrid crops, such as maize or sunflower, is to model the hybrid vigor in the linear mixed models, considering the relatedness between individuals. Here, we compared two additive and three non-additive association models for their ability to identify genomic regions associated with flowering time in sunflower hybrids. A panel of 452 sunflower hybrids, corresponding to incomplete crossing between 36 male lines and 36 female lines, was phenotyped in five environments and genotyped for 2,204,423 SNPs. Intra-locus effects were estimated in multi-locus models to detect genomic regions associated with flowering time using the different models. Thirteen quantitative trait loci were identified in total, two with both model categories and one with only non-additive models. A quantitative trait loci on LG09, detected by both the additive and non-additive models, is located near a GAI homolog and is presented in detail. Overall, this study shows the added value of non-additive modeling of allelic effects for identifying genomic regions that control traits of interest and that could participate in the heterosis observed in hybrids.

## 1 Introduction

Currently, several tools are available to geneticists and breeders to identify the genetic control of traits of interest and to improve the performance of animals and plants. A powerful tool for mapping the genes controlling complex traits, association genetics essentially evaluates statistical correlations between the alleles at a given locus and the observed phenotype (Ersoz et al, 2007). Genome-wide association studies (GWAS) have been widely used in the genetics of humans, animals, and plants (Kang et al, 2008, Wang et al, 2016, Yu et al, 2006, Zhang et al, 2010, Zhou et al, 2012). The method was first applied to human genetics (Corder et al, 1993), and the first association study on agronomic data was conducted in 2001 (Thornsberry et al, 2001) in maize with regard to flowering time.

Flowering time (FT) is a key trait in plant biology. Its evolution has been crucial for the domestication of many crop species and their dissemination into new climatic regions (Blümel et al, 2015, Colledge and Conolly, 2007, Izawa, 2007). It is highly heritable, and the gene regulatory network controlling flowering time is very well described, making it an excellent trait to combine quantitative genetics and functional genomics. The impact of environmental cues on flowering time is well documented in the model plant *Arabidopsis thaliana* where a study (Li et al, 2010) identified SNPs that can explain up to 45% of the phenotypic variation of flowering time in a large panel of natural accessions. In sunflower, GWAS are more recent: Fusari et al (2012) on disease resistance, and Nambeesan et al (2015) on branching performed their GWAS with data collected on inbred lines, whereas Cadic et al (2013) studied the genetic control of FT in a panel evaluated in 15 environments as hybrids.

Many crops, such as maize, sunflower and winter oil seed rape, are cultivated as hybrids. Hybrid vigor, or heterosis, was first observed by Darwin (1876). Genetic mechanisms underlying heterosis have been suggested, but their relative importance is not clearly elucidated (Lamkey and Edwards, 1999). Different hypotheses including dominance (Bruce, 1910, Jones, 1917), over-dominance (Crow, 1948), and subsequently epistasis have been proposed (Williams, 1959). Most GWAS models have been designed to consider only the additive effects of markers. Several studies have shown that non-additive effects constitute a major part of the variation of complex traits. These studies consider the intra-locus effects (Gengler et al, 1997, Norris et al, 2010), namely dominance, or inter-locus effects called epistasis (Huang et al, 2012, Mackay, 2014). The work of Yang et al (2014) on corn showed an increase in the proportion of heritability, explained because the model considered the dominance, thus allowing a better overview of heterosis. Mackay (2014) also stated that epistasis might be linked to missing heritability and small additive effects. Before them, Zhou et al (2012) demonstrated on rice hybrids that the accumulation of multiple effects, including dominance and overdominance, might partially explain the genetic basis of heterosis. In human genetics, it has also been shown that models considering non-additive intra-locus effects yield new information, as in the case for the study by He et al (2015), which found three new quantitative trait loci (QTLs) associated with kidney weight, compared to additive models. In contrast, Tsepilov et al (2015) showed in humans that it is preferable to use non-additive effects only for traits where the non-additive function is known. Additive models already capture a small part of the non-additive variability.

Mixed models are among the methods used to perform association analysis. They take into account the dependence between individuals by introducing a covariance structure for the genetic value of each individual and was proposed by Yu et al (2006). The main drawback of the mixed model is its computational burden. So new methods were proposed to accelerate the algorithm speed, EMMA (Kang et al, 2008) that avoid redundant matrix calculation, EMMAX (Kang et al, 2010) that is an approximation method with the ability to handle a large number of markers and finally GEMMA (Zhou and Stephens, 2012) that is exact and efficient. All these methods are based on single-locus tests, but the traits can be controlled by many loci, with broader effects, and these models do not yield a good estimate of the markers effects in this case.

The identification of causal polymorphisms with the adjustment of more than one polymorphism at a time is complicated by the presence of linkage disequilibrium. Several multi-locus approaches have been proposed, including penalized regressions (Hoggart et al, 2008), Lasso (Waldmann et al, 2013, Wang et al, 2011, Yi and Xu, 2008), and even the elastic net (Waldmann et al, 2013). Segura et al (2012) proposed a regression method with inclusion by forward selection. This method involves EMMAX, that reassesses the genetic and residual variances at each step of the algorithm. An assessment of the model quality, based on a selection criterion, is then performed.

The aim of our study was to evaluate different GWAS models that take dominance into account to detect associations in a hybrid panel and patterns of genetic control putatively involved in heterosis. For this purpose, we used the sunflower and flowering time as an example of the genetic control of complex traits, and we performed this study in a variety of environments to introduce realistic phenotypic variability. Several models involving intra-locus non-additive effects that are appropriate for a GWAS were tested. We sought to compare these models and conventional additive models of GWAS based on a multi-locus method similar to the one reported in Segura et al (2012).

## 2 Materials & Methods

### 2.1 Dataset collection

We collected data on the flowering time of sunflower (*Helianthus annuus*) from various French experiments conducted in 2013 by private partners (Biogemma, Caussade Semences, Maisadour Semences, RAGT2n, Soltis, Syngenta France) and by the French National Institute for Agricultural Research (INRA) as part of the SUNRISE project. Five experimental sites in different environments of regions in Southwestern France were planted with different hybrids from a set of 452 hybrids (between 303 to 444 hybrids per environment (Table S1)). Hybrids for this study were obtained by crossing 36 males and 36 females in an incomplete factorial design. They were chosen so that every parent was represented equivalently in the hybrid population (between 12 and 15 hybrids per parent).

In each environment, each measure of flowering time corresponded to one plot, planted with individuals of a single genotype. Each plot varied from 10 to 18*m*^2^ depending on the environment, and the plant density (corresponding to the number of plants per *m*^2^) was 5.8 on average and varied from three to eight plants per *m*^2^. Flowering time was recorded when 50% of the plants in a plot were flowering and was then converted into degree days since the sowing date relative to the base 4.8 °C, using the mean daily air temperature measured at each location.

### 2.2 Genotyping data

SNP genotyping was performed in the same way as submitted in Badouin *et al.*(2017), but here made from an Illumina type assembly. This work allowed us to obtain genotyping data from the 72 parents on 2,204,423 SNPs that were coded depending on the allele that a parent line could transmit to its descendants: 0, 1, or missing (0 for the XRQ allele, 1 for the variant). The genotyping data were imputed by genomic scaffolds by means of BEAGLE (Browning and Browning, 2009). Nevertheless, this step of the imputation of missing data created some redundancy among SNPs. Maintaining the redundancy for further GWAS analyses increases the computational burden. In addition, redundancy included in the calculation of the relatedness between hybrids tends to give more weight to regions containing many redundant markers, decreasing the power in these regions (Rincent, 2014). Redundant SNPs were therefore discarded. One last filter on minor allele frequency (MAF) was implemented. SNPs with MAF (calculated for parent genotypes before imputation) less than 0.1 were discarded. A total of 478,874 non-redundant polymorphic SNPs were finally retained for various subsequent analyses. The genotypic data of hybrids were deduced from the genotypic data of the parents and coded as 0, 1, or 2 for homozygous XRQ and heterozygous and variant homozygous, respectively. In addition, the male and female origin of alleles was recorded for heterozygous SNPs.

### 2.3 Phenotype adjustment

Data were first adjusted using a linear model including two spatial fixed factors (line and column numbers in the field), a replicate fixed factor if necessary, an independent random genetic factor and the residual error.

### 2.4 GWAS

The analyses were performed using a multi-locus approach with forward selection as proposed by (Segura et al, 2012). This method is based on inclusion (at every step) of the SNP with the smallest p-value as a fixed regressor in a model that contains a random polygenic effect, as in classic GWAS model of Yu et al (2006). The polygenic and residual variances are re-evaluated at each step, and a new scan of the remaining genome is performed. The more integrated regressors in the model, the lower the variance attributed to the random polygenic term. The forward selection analysis stops when the proportion of variance explained by this polygenic effect is close to zero.

Five models were compared to find chromosomal regions linked to flowering time.

#### 2.4.1 Two additive models: A_AIS_ and A_XX′_ models

The first model, as described in (Segura et al, 2012), takes into account only the additive effect of markers. Let *y*_*i*_ denotes the adjusted phenotype of hybrid *i*. Then the additive model is

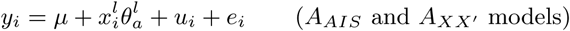

where 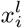 is the centered genotype (coded as XRQ allelic dose) of the *i*th hybrid at the *l*th marker locus; 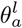 is the additive effect of the *l*th locus; *u*_*i*_ denotes the random polygenic effect; and *e*_*i*_ is the residual error. Let ***u*** and *e* be vectors (*u*_*i*_, *i* = 1, …, *n*) and (*e*_*i*_, *i* = 1, …, *n*), respectively, and then 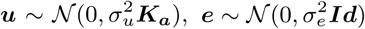, where ***K*_*a*_** is a kinship matrix (relations among hybrids), and 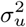 and 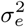 are polygenic and residual variances, respectively.

One simple way to calculate the relatedness between hybrids based on molecular markers is to consider the proportion of shared alleles between two individuals, also called Alike In State (AIS) relatedness.

The formula for biallelic markers is

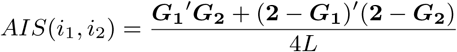

where *L* is the total number of markers, ***G*_1_** and ***G*_2_** are the vector of genotypes for *i*_1_ and *i*_2_ (length of *L*, coded as XRQ allelic dose), and **2** denotes a vector of two. The use of this formula for relatedness between hybrids does not consider haplotypic phases. However, haplotypic phases are known in our factorial design. Accordingly, we consider the AIS between the parents and known haplotypic phases to calculate the relatedness between hybrids. Thus, the AIS kinship that was used in the additive model designated the *A*_*AIS*_ model was calculated as the average *AIS* between respective parents of hybrids.

The other relationship matrix, used in the additive model designated the *A*_*XX*′_ model, is equivalent to the unscaled kinship matrix described by Vanraden (2008):

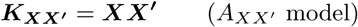

where 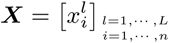 is the centered matrix of the hybrid genotypes.

#### 2.4.2 The additive and dominant model: AD model

A model including additive and dominant effects of SNP markers as proposed by Su et al (2012) was studied next. The model is

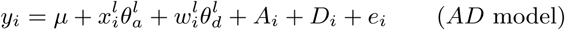

where 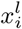 is the centered genotype of the *i*th hybrid at the *l*th marker locus; 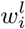 is defined later; 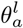 is the additive effect of the *l*th locus; 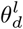 is the dominance effect of the *l*th locus; and *e*_*i*_ denotes error. *A*_*i*_ is the random additive effect *i*, and *D*_*i*_ is the random dominant effect *i*. Let ***A***, ***D***, and *e* denote vectors (*A*_*i*_, *i* = 1, …, *n*), (*D*_*i*_, *i* = 1, …, *n*), and (*e*_*i*_, *i* = 1, …, *n*), respectively, and then 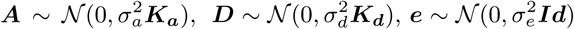, where ***K*_*a*_** is the additive kinship matrix; ***K*_*d*_** is the dominance kinship matrix; and 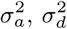 and 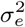 are additive, dominance and residual variances, respectively. ***K*_*a*_** = ***K*′_*XX*_** as in the *A*_*XX*′_ model, and ***K*_*d*_**, = ***WW*′** where 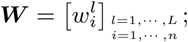; *L* is the number of loci; *n* denotes the number of hybrids; and

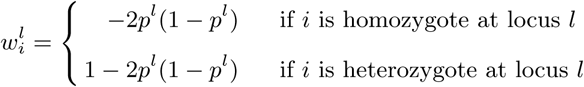

where *p*^*l*^ is the XRQ allelic frequency at locus *l* within the parental population that is equal to the XRQ allelic frequency at locus *l* within the hybrid population under Hardy-Weinberg assumptions.

The part of additive variance used in the forward selection algorithm as a stopping criterion was defined in MLMM (Segura et al, 2012) by 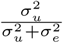. To generalize the stopping criteria for the *AD* model, we used the ratio 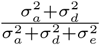.

#### 2.4.3 The models with female and male effects: FM and FMI model

These models include the male and female effects of SNP markers. The last also includes the interaction between male and female effect. Let *y*_*ƒm*_ denote the adjusted phenotype of hybrid obtained when female line *ƒ* was crossed with male line *m*, and then the model is

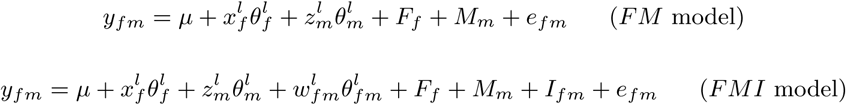

where 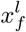 is the centered (0 or 1) allele transmitted by female *ƒ* at the *l*th marker locus; 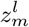 is the centered (0 or 1) allele transmitted by male *m* at the *l*th marker locus; 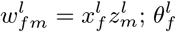 is the female effect of the *l*th locus; 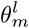 is the male effect of the *l*th locus; and 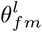 is the female-male interaction effect of the *l*th locus. *F*_*ƒ*_, *M*_*m*_, and *I*_*ƒm*_ are the random effects of female *ƒ*, male *m*, and their interaction, respectively, and *e*_*ƒm*_ denotes error. Let ***F***, ***M***, ***I***, and *e* denote vectors (*F*_*ƒ*_, *ƒ* = 1, …, *n*_*ƒ*_), (*M*_*m*_, *m* = 1, …, *n*_*m*_), (*I*_*ƒm*_, *ƒ* = 1, …, *n*_*ƒ*_; *m* = 1, …, *n*_*m*_), and (*e*_*ƒm*_, *ƒ* = 1, …, *n*_*ƒ*_; *m* = 1, …, *n*_*m*_), respectively, where *n*_*ƒ*_ and *n*_*m*_ are the numbers of females and males, respectively. 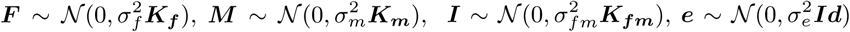 where ***K*_*ƒ*_** is the kinship matrix for the female; ***K*_*m*_** is the kinship matrix for the male; ***K*_*ƒm*_** is the kinship matrix for the interaction between the male and female; and 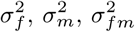 and 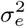 are the female, male, female by male interaction and residual variances, respectively. 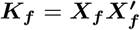 and 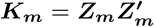 as in the *A*_*XX*′_ model but now using the centered matrix of transmitted alleles, and 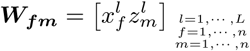 is the Hadamard product between ***X*_*ƒ*_** and ***Z*_*m*_**.

The stopping criterion of the algorithm was dened by the ratio 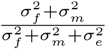 and 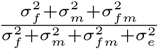 or the *FM* and the *FMI* model, respectively.

#### 2.4.4 Model selection and detected SNP estimation

The main problem of the multi-locus analysis is how much to integrate the SNPs into the model. BIC (Bayesian Information Criterion), which is generally used, is not strict enough for model selection in large model space (Chen and Chen, 2008). Accordingly, eBIC (extended Bayesian Information Criterion), an extension of BIC, was developed (Chen and Chen, 2008). It penalizes the BIC calculation by taking into account the number of possible models for a given number of regressors in the model using mathematical combination, also known as the binomial coefficient. For our models, the total and the given numbers of regressors used in mathematical combination depend on the SNP numbers and SNP modeling and are as follows:

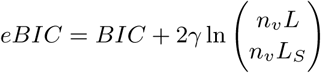

where *L* is the total number of SNPs; *n*_*v*_ is the number of variance components other than residual variance in the model; *L*_*s*_ is the given number of SNPs in the model; 0 ≤ γ ≤ 1 and 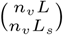 is the mathematical combination of *n*_*v*_*L*_*s*_ among *n*_*v*_*L*.

One way to choose the best γ is to find *k* so that *L* = *n*^*k*^ and then to assume 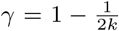 (Chen and Chen, 2008).

To calculate the effects of SNPs selected by eBIC, the model *FMI*, which is the most complete model, was used. It was composed of all eBIC-selected SNPs. Tukeys test of mean comparison was then performed to analyze the significance of differences among the four genotypic classes (00, 01, 10, and 1 1).

### 2.5 Linkage disequilibrium

Linkage disequilibrium was studied to compare and pool the discovered SNPs among models and environments. It was calculated between all pairs of SNPs selected by eBIC, using the classic *r*^2^ (squared Pearson’s correlation) of the hybrid parent genotypes (i.e., SNP correlation of 36 males and 36 females). The significance level of linkage disequilibrium was found by randomly sampling independent SNPs. A total of 10,000 random pairs of SNPs (from 478,874) belonging to different chromosomes were processed. The significance threshold was computed as the 99% quantile of the 10,000 *r*^2^ distribution. We therefore focused on linkage disequilibrium values higher than this threshold.

### 2.6 QTL definition

The use of QTLs instead of SNPs allows us to identify regions of interest rather than specific loci. A QTL is defined as a group of SNPs located on the same chromosome with linkage disequilibrium greater than the predefined significance threshold, or an isolated SNP associated with a trait without the above characteristics. Since the 13EX03 and 13EX04 environments were not properly randomized, isolated SNPs from these environments were removed from the study.

For functional analysis, one SNP per QTL was selected as representative of the QTL. This choice was made based on the test p-value in a SNP by SNP model *FMI*. If a given SNP was associated with a trait in several environments, one p-value per environment was calculated, and the minimal p-value was assigned to the SNP. The SNP ultimately representing the QTL is the one with the lowest p-value.

## 3 Results

### 3.1 Phenotypic data analysis

The period from sowing date to flowering time was measured in various environments. The flowering time in each environment was assumed to be a separate trait. Genotypic variance differed significantly from zero in all environments. The proportion of variance explained by genotypes (usually defined as broad sense heritability) ranged from 0.78 to 0.94 (Table 1).

**Table 1:** Proportion of variance explained by genotypes (*h*^2^) per environment (13EX01 to 13EX06)

|  | 13EX01 | 13EX02 | 13EX03 | 13EX04 | 13EX06 |
| --- | --- | --- | --- | --- | --- |
| $h^2$ | 0.86 | 0.79 | 0.94 | 0.91 | 0.88 |

The correlations among environments are high (Figure S1), ranging from 0.68 to 0.85. Environment 13EX02 correlates with the others the least, with correlation coefficients between 0.679 and 0.697. This result can be explained by the fact that the sowing date for this environment was 7 to 20 days after the other sowing dates. In addition, measurement in this environment was performed less regularly. Despite the good correlation among environments, we analyzed each one independently to capture environment-specific associations.

### 3.2 SNPs associated with the trait

Table 2 shows the number of associated SNPs in each model by environment. For analysis involving models that consider only additive effects (*A*_*AIS*_ and *A*_*XX*′_), the number of associated SNPs ranges between two for model *A*_*AIS*_ in environment 13EX01, for example, and eight for the same model in environment 13EX03. In the analysis with model *A*_*AIS*_, the number of SNPs associated with the trait is greater or equal to the number of SNPs in model *A*_*XX*′_ for all environments except 13EX01. For association analysis involving models other than additive ones (*AD*, *FM*, and *FMI*), the eBIC selection only retains a single SNP.

**Table 2:** Number of SNPs associated with flowering time selected by the forward approach and eBIC per environment and per model. The results for additive models with different kinships *A*_*AIS*_ and *A*_*XX*′_) and non-additive models including dominance (*AD*), female and male effects (*FM*), and female, male and their interaction effects (*FMI*), are presented in five environments (13EX01 to 13EX06).

|  | 13EX01 | 13EX02 | 13EX03 | 13EX04 | 13EX06 |
| --- | --- | --- | --- | --- | --- |
| $A_{AIS}$ | 2 | 3 | 8 | 4 | 6 |
| $A_{XX'}$ | 4 | 3 | 5 | 4 | 2 |
| $AD$ | 1 | 1 | 1 | 1 | 1 |
| $FM$ | 1 | 1 | 1 | 1 | 1 |
| $FMI$ | 1 | 1 | 1 | 1 | 1 |

The MLMM approach selects a single SNP, i.e., the most associated one, to explain the effect of the causal polymorphism in this genomic region. However, several SNPs could be in LD with the causal polymorphism and different sources of errors (phenotypic and genotypic), and missing data could lead to the selection of different SNPs to explain the same causal polymorphism in our different experiments. Therefore, we grouped the SNPs to define QTLs and refer to regions rather than specific positions. This grouping was achieved using linkage disequilibrium between SNPs and positions on the sunflower genomic reference sequence (Badouin *et al*, 2017).

### 3.3 Estimation of linkage disequilibrium (LD)

All SNP pairs with *r*^2^ (squared Pearson’s correlation) above 0.155 were considered to be in linkage disequilibrium. This significance threshold was defined as the 99% quantile of the *r*^2^ distribution obtained for 10,000 randomly sampled pairs of independent SNPs.

We studied the linkage disequilibrium between the SNPs selected by eBIC for all models and environments. Figure 1 illustrates (according to the physical positions of SNPs in the reference genome) only disequilibria greater than the significance threshold of 0.155. Pairs of SNPs located on chromosome LG01, LG11 and LG16 are in strong LD. A LD block is located on chromosome LG09 (*r*^2^ between 0.29 and 0.93). One SNP in disequilibrium with this group is itself located on chromosome LG07. These LDs correspond either to long-range disequilibria that can be caused by imperfect positioning of contigs in the reference genome or to the limited size of our parental population. With the statistical risk at 1% (it should be reduced to take into account the multiplicity of LD tests between all pairs of discovered SNPs), we obtained a threshold of 0.155, which is slightly lower than the linkage disequilibrium thresholds used in other association studies on the sunflower (*r*^2^ = 0.2 reported by Cadic et al (2013) and Nambeesan et al (2015)). In total, this approach allowed us to build 13 associated regions (QTLs) for flowering time on 11 chromosomes.

**Fig. 1:**
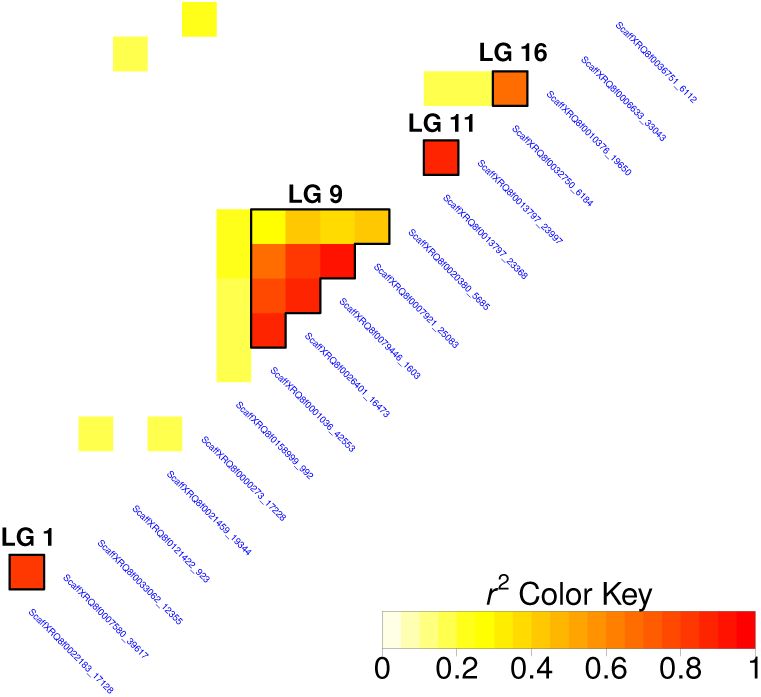
Heatmap of linkage disequilibria between SNPs associated with the flowering time, among all environments and models. Only linkage disequilibria above the significance threshold of 0.155 was represented. Black lines highlight linkage disequilibria between SNPs on the same chromosome. The linkage group (LG) is indicated above a group of interest in black.

### 3.4 QTL description

Groups of five or two SNPs in LD together with single SNPs define the QTLs presented in Table 3. It is noteworthy that four of the five SNPs defining QTL FT09.199 were obtained with non-additive association models. Similarly, the FT11.47 region was only detected by non-additive models. The FT15.102 region was detected by the model taking into account male and female effects and by both additive models in all environments (Table S2). The last eight QTLs were detected by additive models only and tend to have higher p-values than non-additive QTLs.

**Table 3:** List of QTLs associated with flowering time. For each QTL, the following information on the detected SNP is presented: chromosome (LG), position (bp), minor allele frequency (MAF), GWAS model: additive with different kinships *A*_*AIS*_ and *A*_*XX*′_) and non-additive including dominance (*AD*), female and male effects (*FM*), and female, male and their interaction effects (*FMI*), and p-values calculated in the *FMI* model, incorporating only the detected SNP. For each QTL composed of several SNPs, the SNP with the smallest p-value is highlighted in bold.

| QTL | SNP | LG | Position | MAF | Models | p-value |
| --- | --- | --- | --- | --- | --- | --- |
| FT09.199 | ScaffXRQ8f0001036_42553 | 9 | 198,931,169 | 0.26 | $AD, FMI$ | $1.84 \times 10^{-11}$ |
| | **ScaffXRQ8f0026401_16473** | 9 | 199,047,735 | 0.32 | $AD, FMI, FM$ | $8.67 \times 10^{-13}$ |
| | ScaffXRQ8f0079446_1603 | 9 | 199,131,966 | 0.33 | $AD, FMI$ | $1.86 \times 10^{-08}$ |
| | ScaffXRQ8f0007921_25083 | 9 | 199,145,681 | 0.29 | $A_{XX'}$ | $6.14 \times 10^{-09}$ |
| | ScaffXRQ8f0020380_5685 | 9 | 201,493,137 | 0.24 | $AD, FMI$ | $3.57 \times 10^{-09}$ |
| FT11.47 | ScaffXRQ8f0013797_23368 | 11 | 47,534,503 | 0.42 | $AD$ | $4.16 \times 10^{-07}$ |
| | **ScaffXRQ8f0013797_23997** | 11 | 47,535,132 | 0.39 | $FM, FMI$ | $3.62 \times 10^{-07}$ |
| FT16.167 | **ScaffXRQ8f0010376_19650** | 16 | 167,723,083 | 0.39 | $A_{XX'}, A_{AIS}$ | $2.76 \times 10^{-04}$ |
| | ScaffXRQ8f0032750_6184 | 16 | 167,689,531 | 0.42 | $A_{XX'}$ | $6.74 \times 10^{-02}$ |
| FT01.98 | **ScaffXRQ8f0007580_39617** | 1 | 98,035,404 | 0.17 | $A_{XX'}, A_{AIS}$ | $6.77 \times 10^{-06}$ |
| | ScaffXRQ8f0022183_17128 | 1 | 91,634,676 | 0.21 | $A_{AIS}$ | $2.40 \times 10^{-04}$ |
| FT15.102 | ScaffXRQ8f0000770_77572 | 15 | 102,863,872 | 0.26 | $A_{XX'}, A_{AIS}, FM$ | $1.91 \times 10^{-06}$ |
| FT02.78 | ScaffXRQ8f0070840_1738 | 2 | 78,884,560 | 0.11 | $A_{XX'}, A_{AIS}$ | $3.55 \times 10^{-06}$ |
| FT17.184 | ScaffXRQ8f0036751_6112 | 17 | 184,825,665 | 0.18 | $A_{XX'}, A_{AIS}$ | $4.64 \times 10^{-03}$ |
| FT05.208 | ScaffXRQ8f0006894_28213 | 5 | 208,225,977 | 0.21 | $A_{AIS}$ | $1.42 \times 10^{-01}$ |
| FT04.144 | ScaffXRQ8f0065196_696 | 4 | 144,357,532 | 0.36 | $A_{AIS}$ | $1.34 \times 10^{-04}$ |
| FT07.34 | ScaffXRQ8f0001757_13384 | 7 | 34,580,910 | 0.11 | $A_{AIS}$ | $4.15 \times 10^{-02}$ |
| FT17.13 | ScaffXRQ8f0006633_33043 | 17 | 13,852,550 | 0.39 | $A_{XX'}$ | $3.10 \times 10^{-02}$ |
| FT04.74 | ScaffXRQ8f0021459_19344 | 4 | 74,011,326 | 0.19 | $A_{XX'}$ | $1.65 \times 10^{-03}$ |
| FT13.190 | ScaffXRQ8f0023382_14615 | 13 | 190,953,163 | 0.12 | $A_{XX'}$ | $8.91 \times 10^{-01}$ |

The number of QTLs in common within a model and among environments is presented in Table 4. For each model, Table 4 shows the number of QTLs associated in several environments. Five (FT16.167, FT17.184, FT05.208, FT04.144, and FT07.34) and six QTLs (FT09.199, FT01.98, FT17.184, FT17.13, FT04.74, and FT13.190) were associated in only one environment using additive models *A*_*AIS*_ and *A*_*XX*′_, respectively. In contrast, FT15.102 was associated in all five environments for these two models. For non-additive models (*AD*, *FM*, and *FMI*), no QTL appeared in the five environments. Models *AD* and *FMI* detected a QTL in a single environment (FT11.47) and a QTL in four environments (FT09.199). The *FM* model identified two QTLs (FT15.102 and FT11.47) in a unique environment and FT09.199 in two environments.

**Table 4:** Number of QTLs in common within a model and among environment. The results for additive models with different kinships *A*_*AIS*_ and *A*_*XX*′_) and non-additive models including dominance (*AD*), female and male effects (*FM*), and female, male and their interaction effects (*FMI*) are presented for different numbers of environments (env).

| | $A_{AIS}$ | $A_{XX'}$ | $AD$ | $FM$ | $FMI$ |
| --- | --- | --- | --- | --- | --- |
| 1 env | 5 | 6 | 1 | 2 | 1 |
| 2 env | 2 | 2 | - | 1 | - |
| 4 env | - | - | 1 | - | 1 |
| 5 env | 1 | 1 | - | - | - |

### 3.5 QTL effects

We characterized the effects of the SNPs detected in both additive and non-additive models. Regarding QTLs detected by the additive models, the majority of SNPs have a clearly additive profile similar to Figure 2a. However, for some additive SNPs, Tukey’s mean comparison test did not separate the genotypes in three significantly different classes certainly because of a lack of power.

**Fig. 2:**
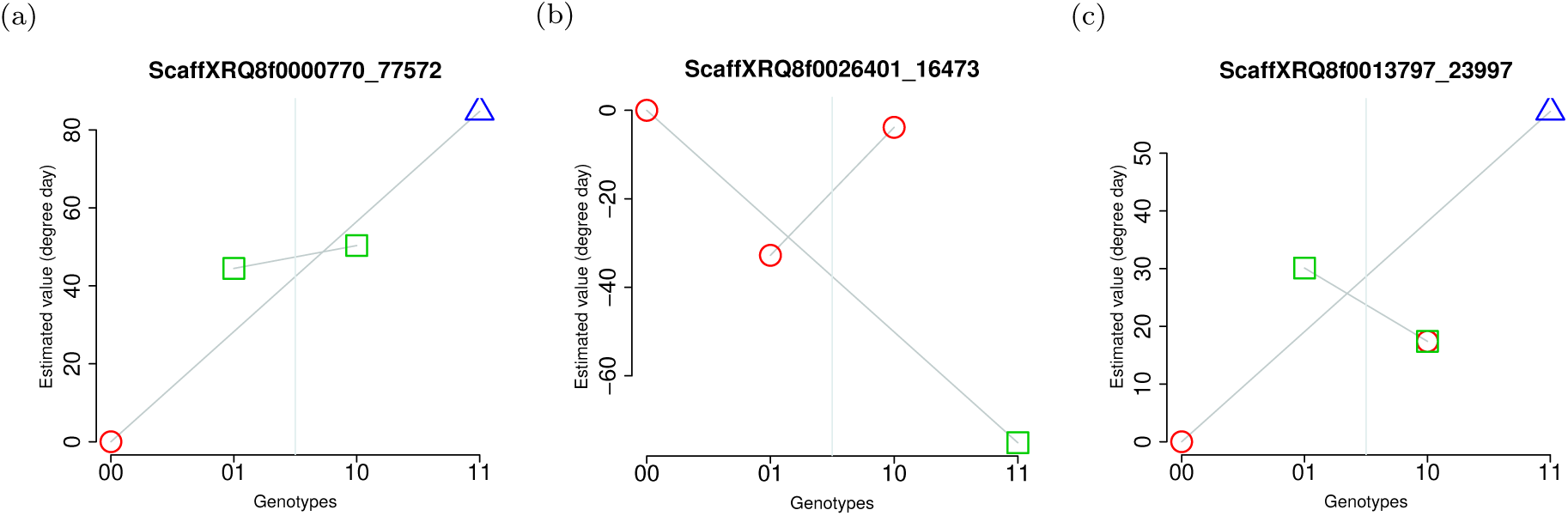
Effects of SNPs on flowering time for the four genotypic classes. (a) Example of an additive SNP. (b) SNP discovered with non-additive model and with an additive trend. 00 and 11 correspond to homozygous genotypes, 10 to the heterozygous genotype that received allele 1 from the female parent and 01 to the heterozygous genotype that received allele 1 from the male parent. Each symbol indicates membership in a specific class in Tukeys mean comparison test with a 5% statistical risk.

The majority of QTLs detected using non-additive models have a profile similar to Figure 2b, with a dominant trend for one allele (reference allele of inbred line XRQ for the male in the example). Two significantly different classes in the mean comparison test, separating one homozygous genotype from the other genotypes, is expected for a dominant allele. Figure 2c illustrates SNP profiles that are more difficult to interpret. Such profiles could be due to slight dominance of the XRQ allele in males or more probably to an additive SNP and insufficient power of Tukey’s test.

### 3.6 QTL annotations

For each QTL, the SNP with the lowest p-value in the model *FMI* was selected to represent the region. All redundant SNPs were excluded from the GWAS analysis, but in terms of the functionality of the gene, information on the location of redundant SNPs is important. SNPs redundant with SNPs that are representative of a QTL were therefore recovered and analyzed in the same way as other SNPs. The results of this analysis are presented in Table 5. All SNPs redundant with the referent SNP of FT09.199 are also located on chromosome LG09 at positions very close to each other (within a 61 kb interval). Two genes are present in this region, but none is known to be involved in flowering. Four QTLs are also located in the identified genes on chromosomes LG05, LG13, LG16, and LG17. These genes do not correspond to a flowering-related gene. One SNP located on chromosome LG17 is redundant with the referent SNP of FT11.47 and another SNP also on chromosome LG11. This situation may be due to the imperfect quality of the genome.

**Table 5:**
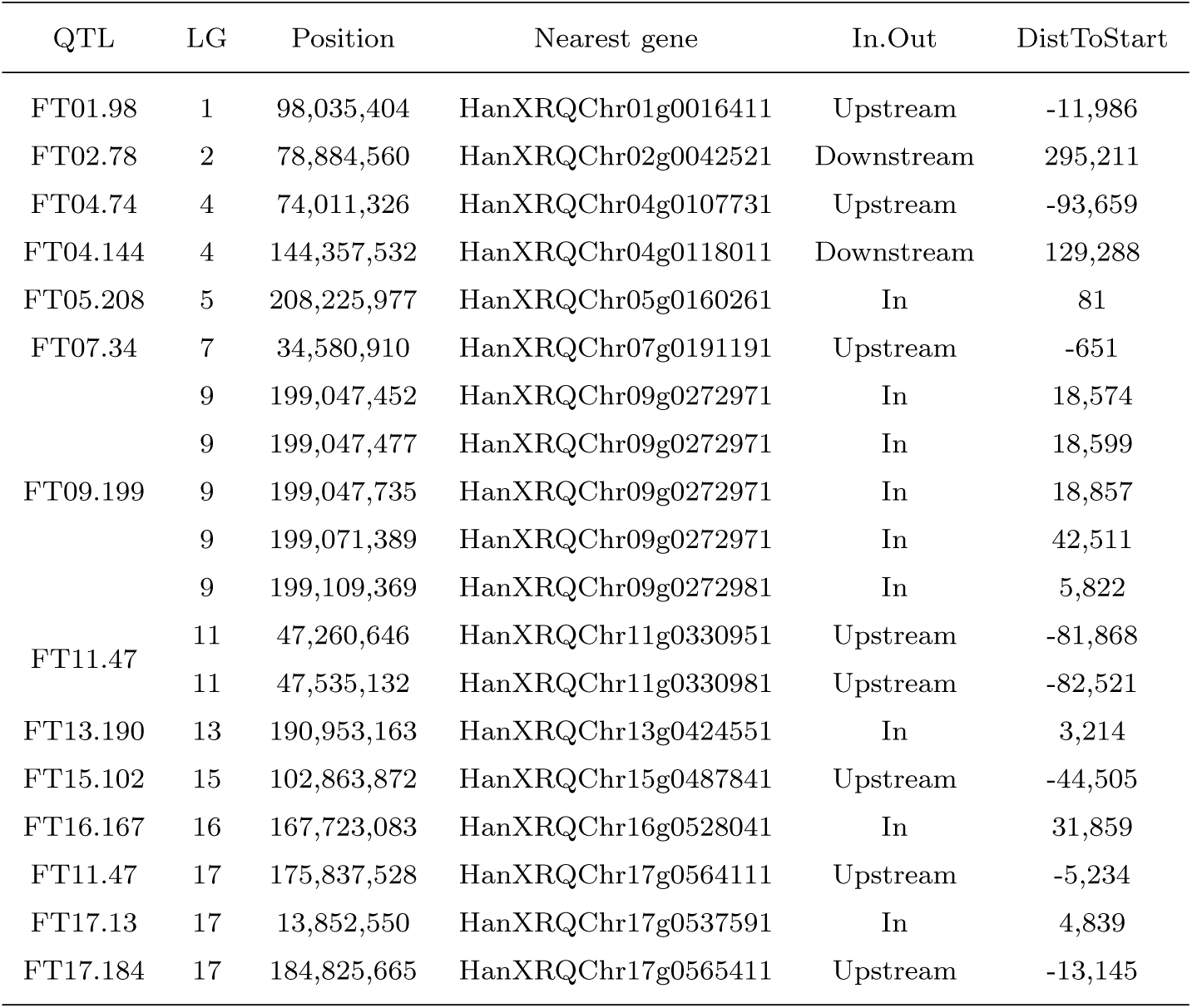
Genes underlying QTLs associated with flowering time. One SNP per QTL was selected, and its redundancy, if applicable, was also analyzed. The table describes QTL name (QTL), chromosome (LG), position (Position), closest gene, location with respect to the closest gene (In.Out) and distance to the start of the closest gene (DistToStart).

Few of the SNPs are located in genes, but three genes known to be involved in the flowering process are located on chromosome LG09. Figure 3 presents the positions of the associated markers and these three genes in FT09.199. *GIBBERELLIC ACID INSENSITIVE* (*GAI*, homologous to *HanXRQChr09g0272901*) is a gene involved in flowering time (Wilson and Somerville, 1995), whereas *FLORICAULA* (*FLO*, homologous to *HanXRQChr09g0273821*) (Coen et al, 1990) and *CAULIFLOWER* (*CAL*, homologous to *HanXRQChr09g0273361*) (Bowman et al, 1993) are genes involved in flowering development. FT09.199 consists of four SNPs very close together (within a 214kb interval) and a more distant SNP (at 2 Mb down chromosome LG09). The most interesting gene based on its function, namely *GAI*, is the closest of the four SNPs. This region was further examined based on the p-values of all SNPs in it. Figure S2 represents p-values for all SNPs of the FT09.199 region and the three genes involved in the flowering. The presented p-values were calculated in the environment and with the model where the SNP of interest was discovered in association. It can be seen that the most significant associations are found in the region of the first four SNPs. With the *FMI* model and for the environments 13EX03 and 13EX06, we can see the SNP with low p-values downstream, i.e., between the two *CAL* and *FLO* genes.

**Fig. 3:**
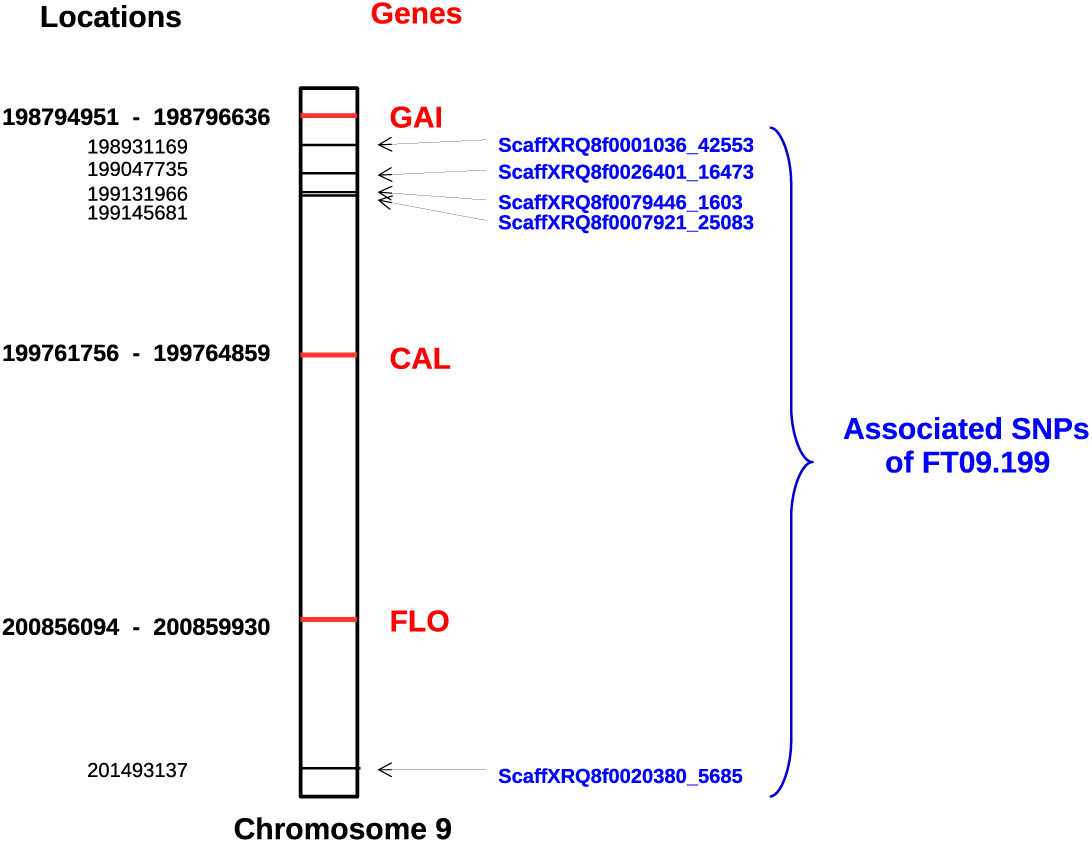
Locations of genes involved in the flowering process, compared to locations of SNPs of FT09.199 located in the same region of the chromosome LG09. Gene and SNP positions are indicated in bold and normal font, respectively. For genes, the two positions correspond to the start and end of the gene.

## 4 Discussion

In this study, we propose new GWAS models including non-additive effects. These models were developed to better model the biological factors involved in sunflower trait variability. Indeed, the modeling of intra-locus effects with a dominance component can capture part of heterosis (Larièpe et al, 2012, Reif et al, 2012), a phenomenon usually observed in sunflower hybrids (Cheres et al, 2000). In addition, the modeling of differences in male and female allelic effects takes into account the two sunflower breeding groups, for which divergence between the maintainer and restorer germplasm has previously been observed by Gentzbittel et al (1994). As in the common additive GWAS model, there is a one-to-one correspondence, in these models between a non-additive fixed effect and its random effect. The rationale of such modeling is to consider multiple QTLs in linkage equilibrium with the currently tested locus and to achieve the same modeling for all the QTLs. Let us suppose that all causal QTLs of the trait genetic architecture are known and located in the genome. A correct model to test each QTL effect while accounting for the other QTLs is a multiple QTL model. Because the QTL locations are unknown, this perfect model is unknown, and a model assuming a QTL effect on each marker is considered. However, the number of parameters in this model is then larger than the number of observed individuals, and to address this issue, it is necessary to abandon the least square estimation method and to use, for example, the L2 shrinkage method. The solution is to assume a normal distribution for the marker effects in linkage equilibrium with the tested locus *l*. Then, as in the equivalence between the rrBLUP method (Endelman, 2011) and the GBLUP method (Vanraden, 2008), it leads to as many fixed effects as random effects, depending on the non-additive model used. The kinship matrices of random effects are proportional to ***XX***′, where ***X*** is linked to the marker effect modeling as in the additive model and the Vanraden kinship matrix (Vanraden, 2008). The computational burden required for computing kinship matrices at each location *l* is then lessened by using the same kinship matrices for all *l*. As shown by Rincent (2014), having different kinship matrices improves the power of GWAS analyses, since it avoids absorbing a part of the signal roughly proportional to linkage disequilibrium in the region of *l* when testing at location *l*.

The difference between the two additive models lies in the kinship matrix computation: one is the usual Vanraden (2008) matrix and the other an AIS-like matrix that takes into account known marker phases in hybrids. Both models detected the greatest number of associated SNPs and of QTLs in common. Indeed, five QTLs are found associated using both additive models and, in particular, FT15.102 was detected in all environments. The QTLs that differ between these models have higher p-values and therefore are less strongly associated with the phenotype. Overall, the two additive models yield coherent results, especially on strongly associated QTLs. Strandén and Christensen (2011) demonstrated that the use of a Vanraden (2008) or AIS relatedness matrix, gives the same prediction of additive genetic values in the GBLUP genomic selection model, and in particular, proved that both matrices give the same REML estimates of random variance components. Therefore, the Wald tests performed in the GWAS forward approach are identical for the two relatedness matrices. Although we used more information in our AIS-like matrix because we integrated the known marker phases, we did not obtain a power improvement in QTL detection, as could be expected.

FT09.199 was found to be associated with flowering time with four models out of five. This region is located on chromosome LG09, and this chromosome was also highlighted by Cadic et al (2013). In their study, the region is found to be associated in six different environments (i.e., combinations Sites × Years). Together, these findings suggest that this QTL is the most interesting region in our study. In addition, 3 genes (*GAI* (Wilson and Somerville, 1995), *FLORICAULA* (Coen et al, 1990) and *CAULIFLOWER* (Bowman et al, 1993)) known to be involved in flower development are also located on chromosome LG09. It is surprising that none of our results falls exactly into these 3 genes, but FT09.199 is near *GAI*. It is likely that the causal polymorphism could be close and in strong linkage disequilibrium with the associated SNPs without being located exactly at the same position. A QTL confidence interval around FT09.199 would be useful to estimate the region where the causal locus should be located. Hayes (2013) proposed a method based on the difference of QTL positions within the region of interest detected in two random subsamples; this method could be applied to our QTL.

GWAS are largely based on additive effect models [58, 44]. Here, three models including non-additive effects were also computed. These models make it possible to emphasize the most interesting region on LG09, as mentioned above. This region is indicated by five SNPs, among which a single SNP was detected by an additive model. The non-additive modeling results increase the reliability of this region through the identification of SNPs very close to the SNP identified using an additive model. Moreover, four SNPs out of five were detected with the *FMI* model, which is the most complex. The usefulness of non-additive models is also illustrated by FT11.47 (on chromosome LG11), since this QTL was only detected with non-additive models despite having a strong impact on flowering time, as illustrated by its effects and p-values in the *FMI* model. In addition, models *AD* and *FMI*, which include intra-locus interaction by modeling dominance or parental allelic interaction, both found the most strongly associated regions indicated by FT09.199 and FT11.47. The first exhibits a clear deviation from an additive behavior, in contrast to the latter, for which an additive behavior cannot be rejected. FT11.47 was not found by additive models, because there is linkage disequilibrium between it and the strong FT16.167 on LG16 detected by additive models. In our forward detection procedure of the additive models, this phenomenon led to the addition of FT16.167 first, which likely decreased the signal of FT11.47 and prevented its detection. Performing GWAS with different models allowed us to increase both the number of associated QTLs and the confidence in the detected regions. Non-additive models can highlight regions with non-additive behavior even for a trait such as flowering time, which is notably genetically additive (Miller et al, 1980, Roath et al, 1982).

However, in our procedure, the extended BIC used to choose non-additive models had two major drawbacks that certainly decreased the number of QTLs detected by these models and thus limited their usefulness. eBIC is an extension of BIC suitable for handling the so-called “high dimension issue” resulting from fewer observations than possible regressors to be put in the model. A penalization term that depends on the number of possible models formed with a given number of regressors is added to BIC in the eBIC calculation (Chen and Chen, 2008). eBIC was established for additive regressor models (Chen and Chen, 2008), and therefore we had to adapt it to the non-additive models *AD, FM*, and *FMI*. The penalization term, which is proportional to the mathematical combination of the number of SNPs in the current model among all SNPs in additive models, was transformed by multiplying both terms of the mathematical combination by the number of SNP effects in non-additive models (2 for *AD* and *FM* and 3 for *FMI*). Nonetheless, it is clear that all possible models are not analyzed during the forward selection process. Indeed, each SNP selected by the algorithm is added to the current model with all its modeling effects. The dominant part of a SNP cannot be added without the additive part, if we take the *AD* model as an example. The number of possible models should have been reduced to take this constraint into account and the penalization term is therefore too high and not completely suitable for non-additive models. Furthermore, the likelihood computation for the eBIC calculation could also be improved. Segura et al (2012) used restricted maximum likelihood (REML) for BIC and eBIC calculations, and we performed our calculations on the same basis. However, REML is not the correct likelihood to compute when selecting a model among models that do not share the same number of fixed effects. This case occurs during the forward selection process, as the algorithm incorporates a new SNP at each iteration. Therefore, we can assume that maximum likelihood (ML) should have been used instead of REML in BIC. Based on this problem, Gurka (2006) performed simulations to compare the use either of REML or ML in model choice criteria. This study demonstrated that REML should incorporate the fixed effects using 1/2 log *det*(***X*′*X***), where ***X*** is the fixed effect design matrix, and showed that this REML approach gave similar or better results than ML in choosing the actual simulated model using BIC. The stringency of eBIC and the absence of a term due to fixed effects may both explain why only a single SNP was selected by the non-additive models for each environment. Nevertheless, as it is more acceptable to exclude too many false negatives than to select too many false positives, we kept the eBIC for our model choice.

Flowering time is an important agronomic trait that impacts crop yield, ecological fitness including adaptation to abiotic factors, and interaction with pollinators. Knowledge of the relative lengths of the period from sowing to flowering is particularly important for breeding yield (Tuteja, 2012), as late hybrids accumulate more biomass than early hybrids, and this advantage can lead a higher yield (Cadic, 2014). For a maximal dry matter yield, all parts of the plant need to develop. This morphology corresponds to late-flowering genotypes (Gallais et al, 1983). The aim of breeders is to find genotypes with the best performance, so regarding the selection of sunflower lines, studies tend to select late lines. Precocity is linked to yield, and therefore the variability of the effect of SNP associated with the flowering time for different genotypes is of interest. The genotypic effects of FT09.199 are far from an additive profile, with homozygotes for the variant allele that flower earlier than the other genotypes. In terms of degree days, 80 degree days separate homozygotes for the variant allele from homozygotes that received XRQ alleles, i.e., a difference of nearly 6 days. This effect is very important regarding the observed variability of approximately 15 days in the multi-environment trials.

The two breeding pools of sunflower (maintainers and restorers of male sterility) have undergone neither the same trait improvement nor the same selection pressure (Mandel et al, 2011). It is therefore expected that modeling different effects for each parental allele, as in *FM* and *FMI* models, will yield different results from other models. However, this expected difference in QTL detection is not obvious in our study, since only a single QTL was detected exclusively by the *FM* and *FMI* models. A lack of differentiation between the two breeding pools in this study compared to Mandel et al (2011) or the too-small number of non-additive QTLs detected because of eBIC could explain this result. Furthermore, even if there are highly differentiated regions between pools, they may not be involved in flowering time variability, as branching and restorer of cytoplasmic male sterility are located on chromosomes LG10 and LG13.

Association analyses were conducted within each environment, as was the phenotypic adjustment to correct the data for local field effects. The differences of observed hybrids and the intra-environment adjustments produced a disturbance in the trait of interest from one analysis to another. Moreover, the sources of trait variation may be different (location, climate, soil, cultural practices, and biotic stress), and these variations may reveal different types of QTLs: generalist QTLs whose action does not depend on the environment and specific QTLs with action that is revealed only in a specific environment. Apart from Cadic et al (2013) in sunflower, this behavior was also observed in *Brassica napus* in Li et al (2015), with a multi-environment GWAS on flowering time. In this study, 44% of all SNPs identified at 3 different locations were found in only one of them. In our study, we detected 6 generalist QTLs revealed by one to five SNPs and found in two to five environments and seven specific QTLs. Naturally, the confidence is greater for generalist QTLs that were detected despite the disturbance of the observed panel, and a region indicated by several SNPs, when they exist, could help to define a confidence region for the underlying causal locus.

Non-additive effects, including dominance or overdominance, have been suggested as underlying heterosis. The modeling of non-additive effects in our models captured part of the heterosis observed in hybrids. This study shows the added value of non-additive modeling of allelic effects, and thus the importance of taking into account heterosis, to identify genomic regions controlling traits of interest for sunflower hybrids.

---

**Fig. S1:**
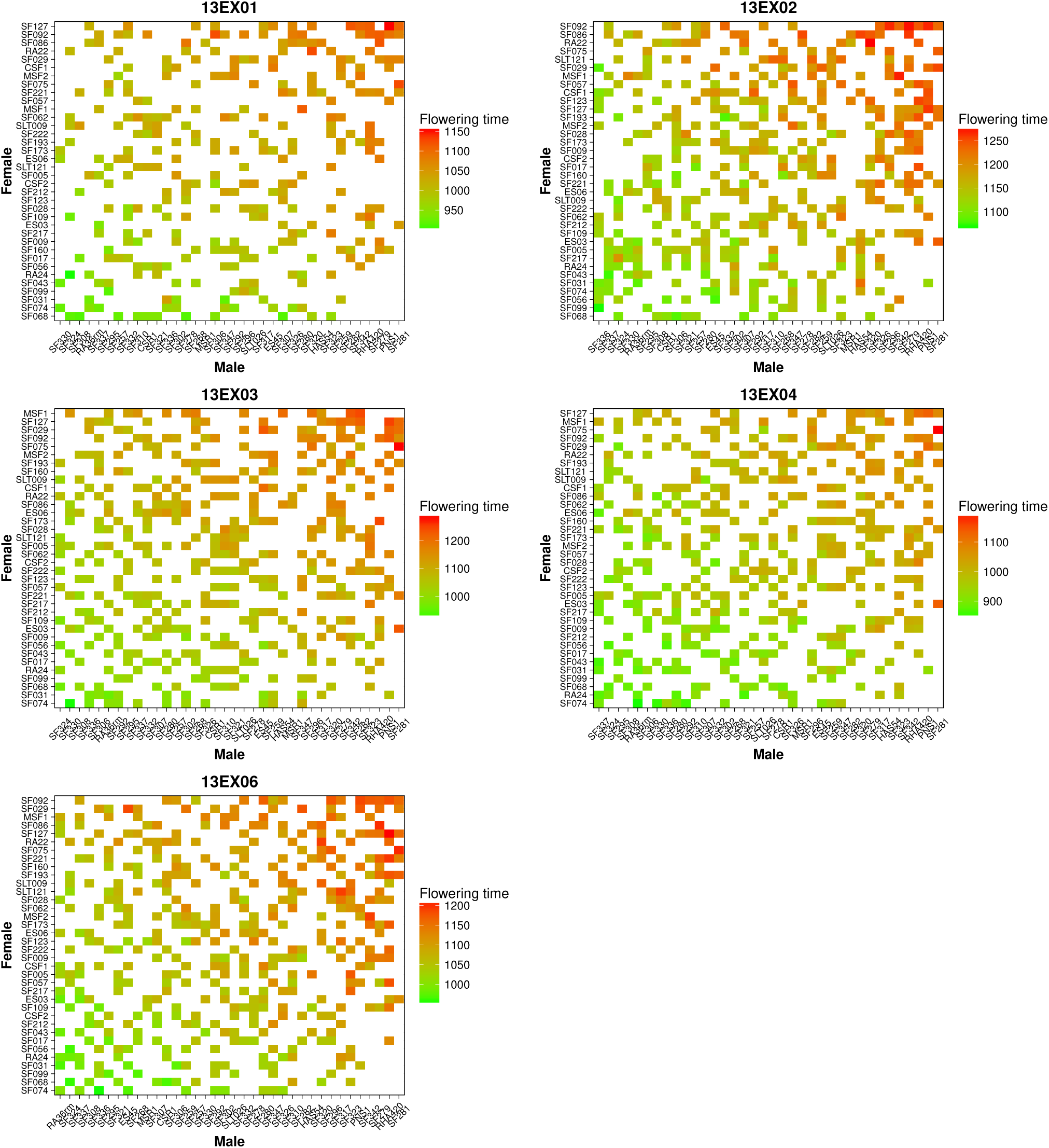
Heatmap of phenotypic values per environment (13EX01 to 13EX06). For each heatmap (one environment), the flowering time values are illustrated by hybrid (one square) according to their parents (female lines on the left and male lines at the bottom). Parents are ranked by the mean of their descendants. The more the square are green, the earlier the hybrid has bloomed, and the more the square are red, more the hybrid has bloomed late.

**Fig. S2:**
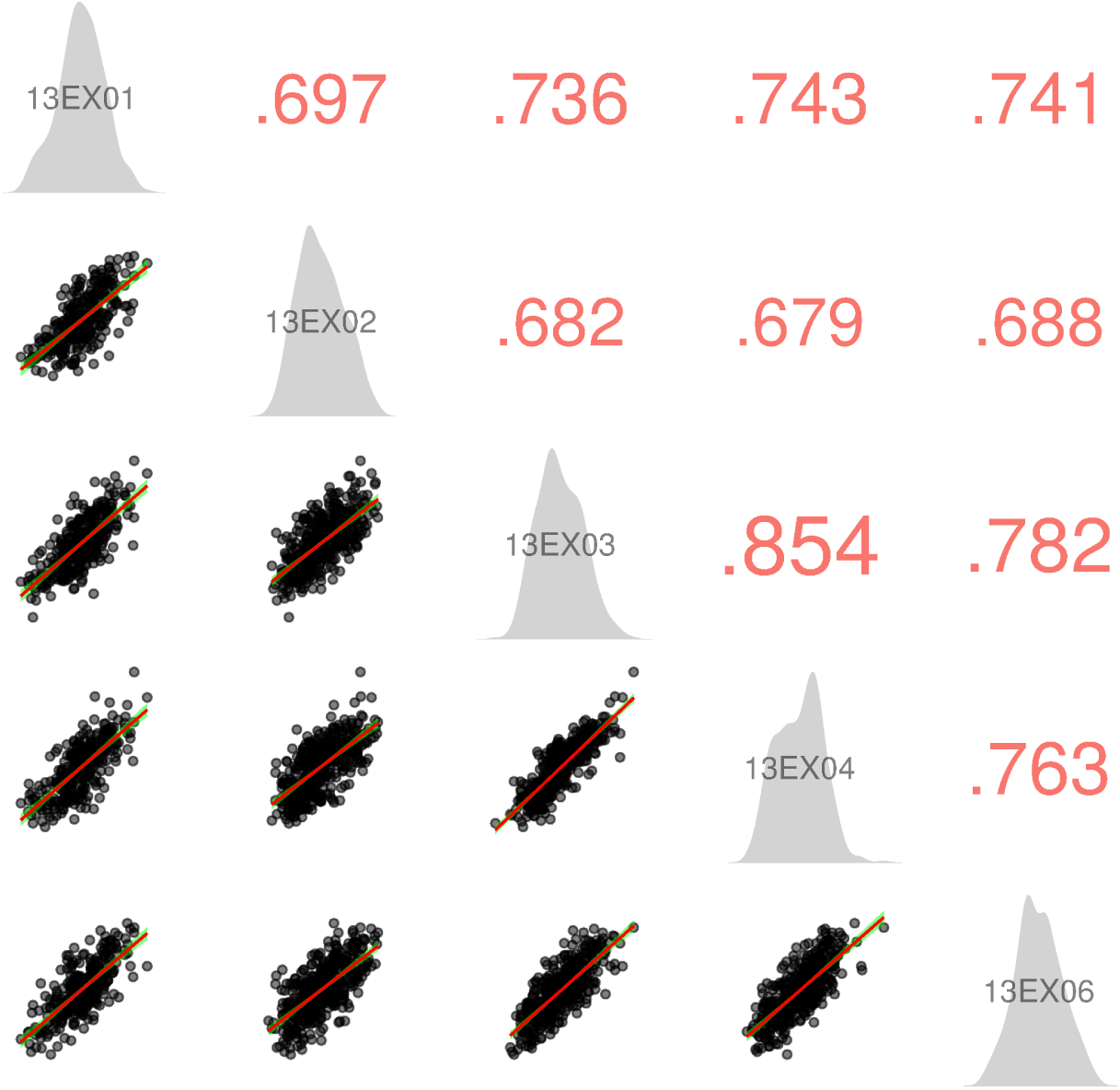
Flowering time in the sunflower hybrid population across environments. Pearsons correlation values are in red on the top right, trait distribution for each environment is presented in gray (middle), and point clouds with the regression slope (red line) with confidence interval (green shading) are presented on the bottom left. Correlations between two environments are based on 370 hybrids on average (minimum 297, maximum 425).

**Fig. S3:**
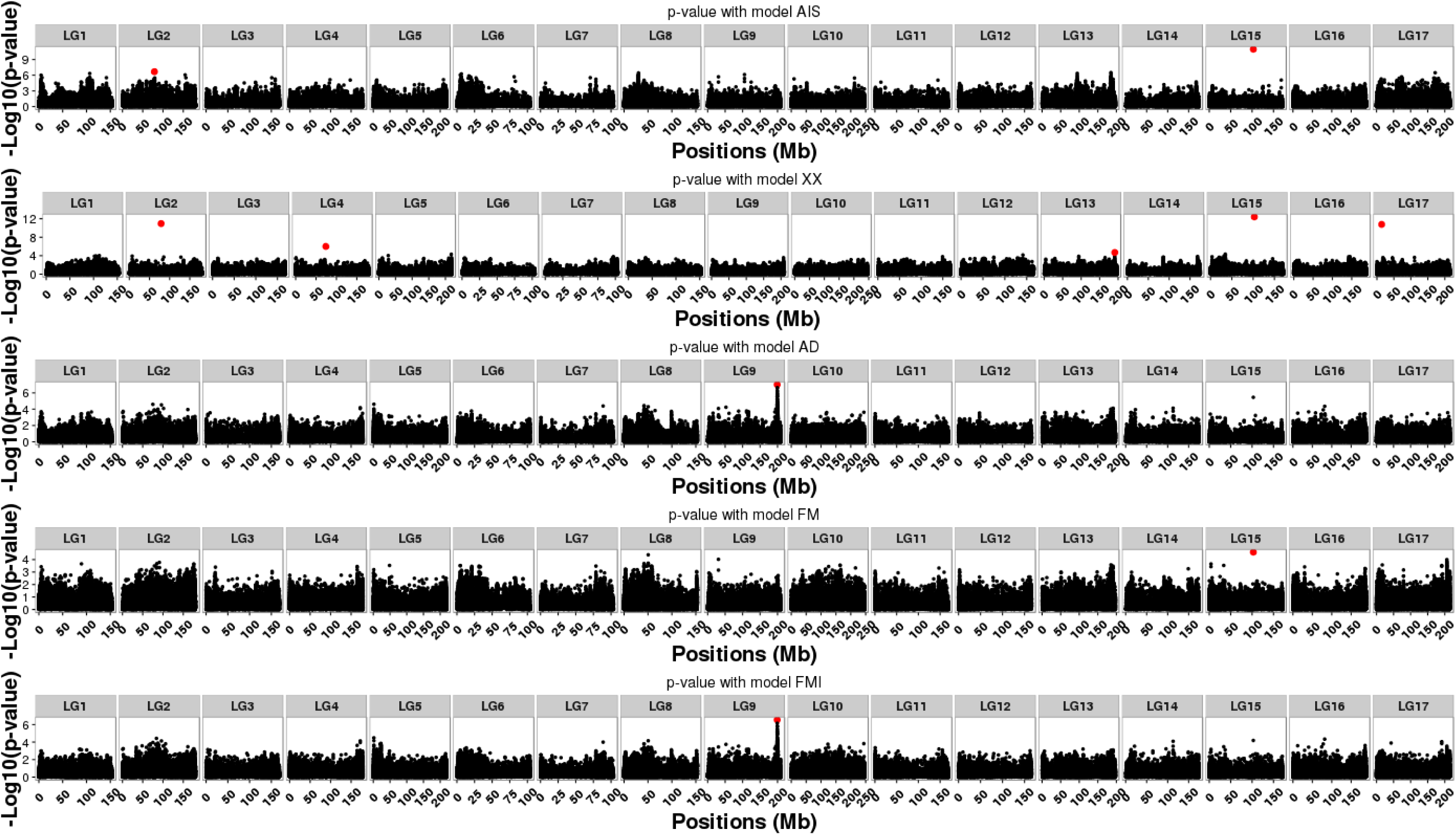
Manhattan plot for all models (*A*_*AIS*_, *A*_*XX*_, *AD*, *FM* and *FMI*) for the environment 13EX01. For each model, the p-values off all SNPs of the analysis are represented according to the position of the SNPs on the genome (in Mb). P-values are those of the step of the algorithm where all SNPs that are part of a QTL (red points) have been detected with the model of interest.

**Fig. S4:**
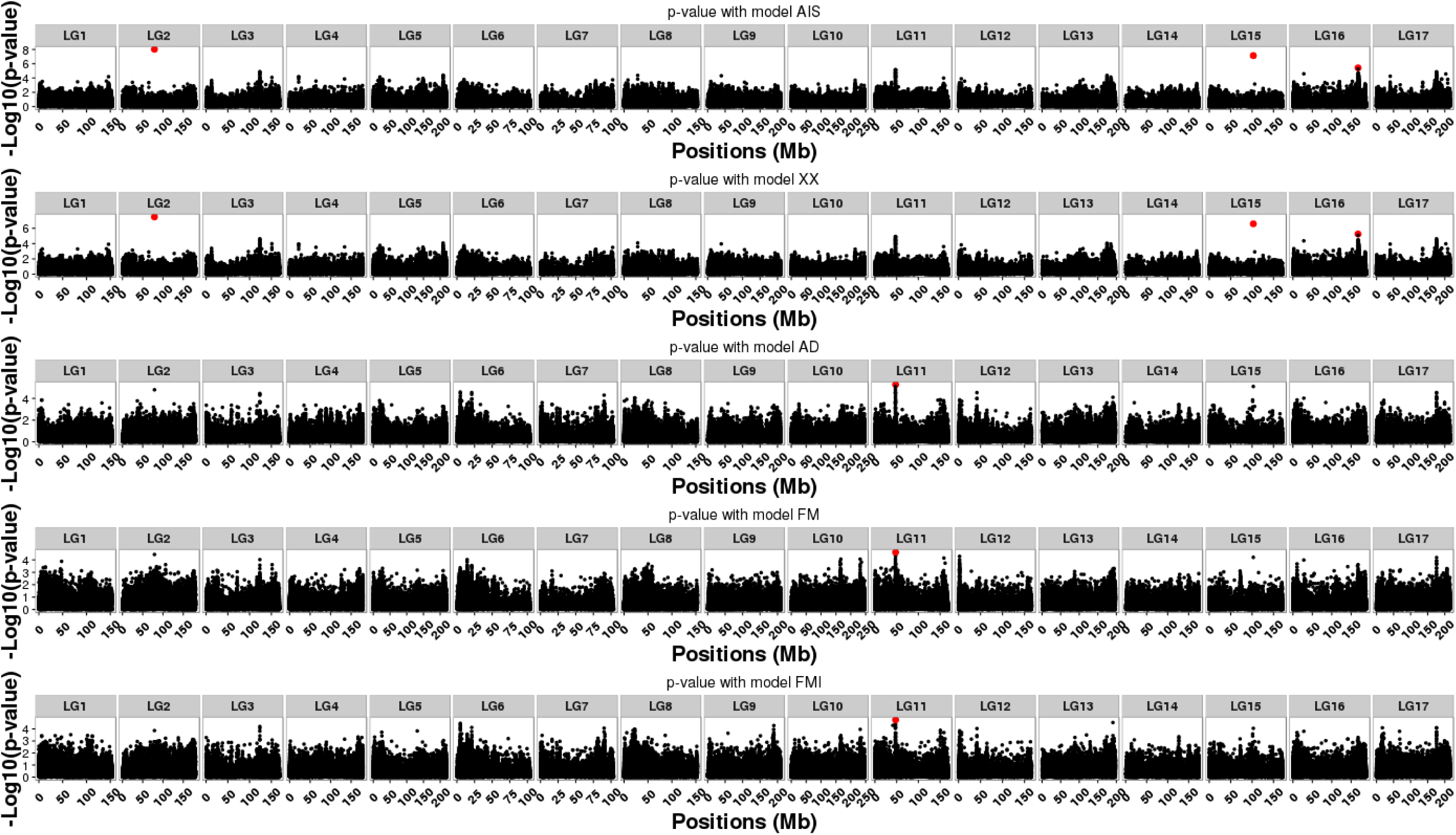
Manhattan plot for all models (*A*_*AIS*_, *A*_*XX*_, *AD*, *FM* and *FMI*) for the environment 13EX02. For each model, the p-values off all SNPs of the analysis are represented according to the position of the SNPs on the genome (in Mb). P-values are those of the step of the algorithm where all SNPs that are part of a QTL (red points) have been detected with the model of interest.

**Fig. S5:**
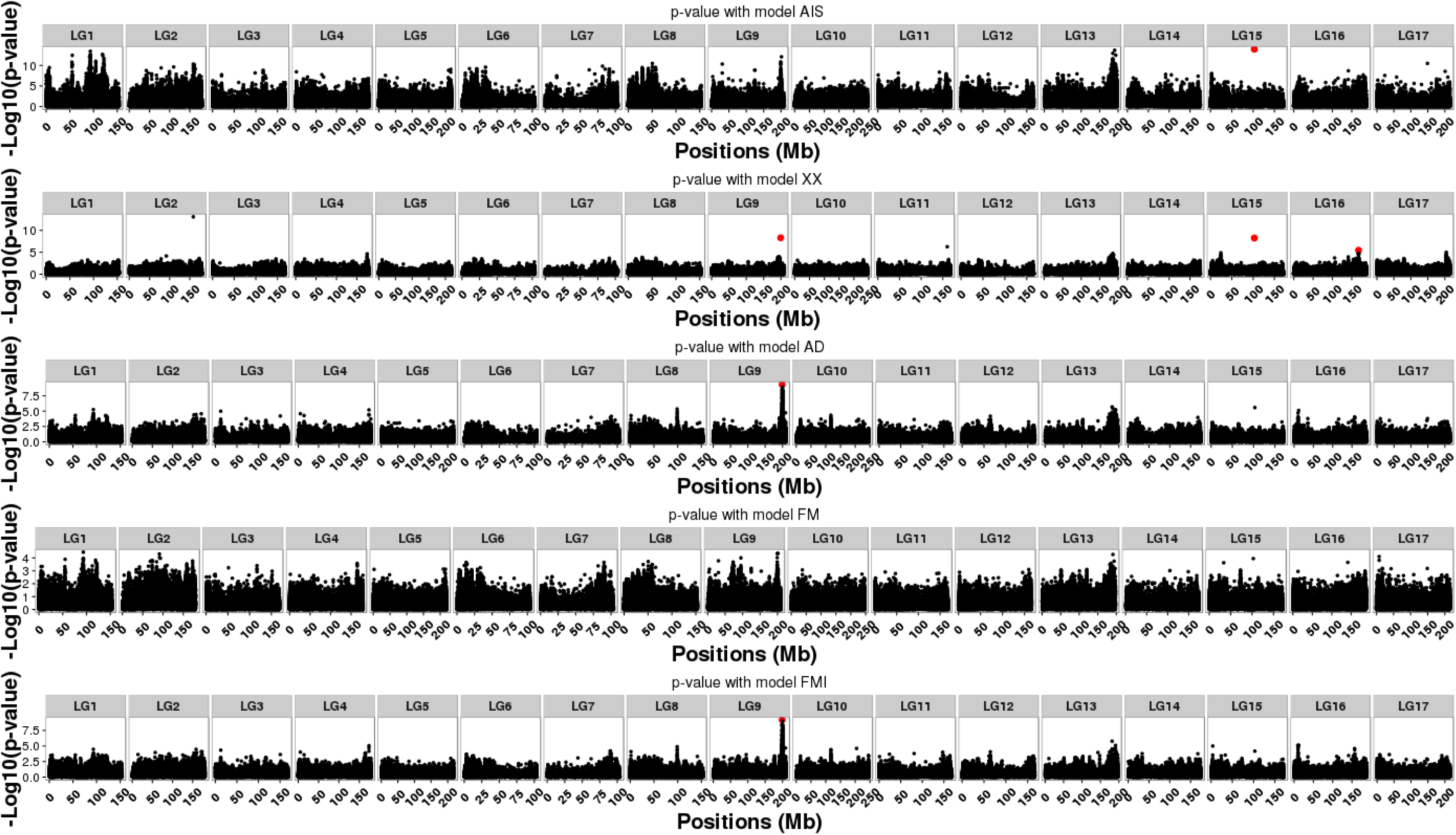
Manhattan plot for all models (*A*_*AIS*_, *A*_*XX*_, *AD*, *FM* and *FMI*) for the environment 13EX03. For each model, the p-values off all SNPs of the analysis are represented according to the position of the SNPs on the genome (in Mb). P-values are those of the step of the algorithm where all SNPs that are part of a QTL (red points) have been detected with the model of interest.

**Fig. S6:**
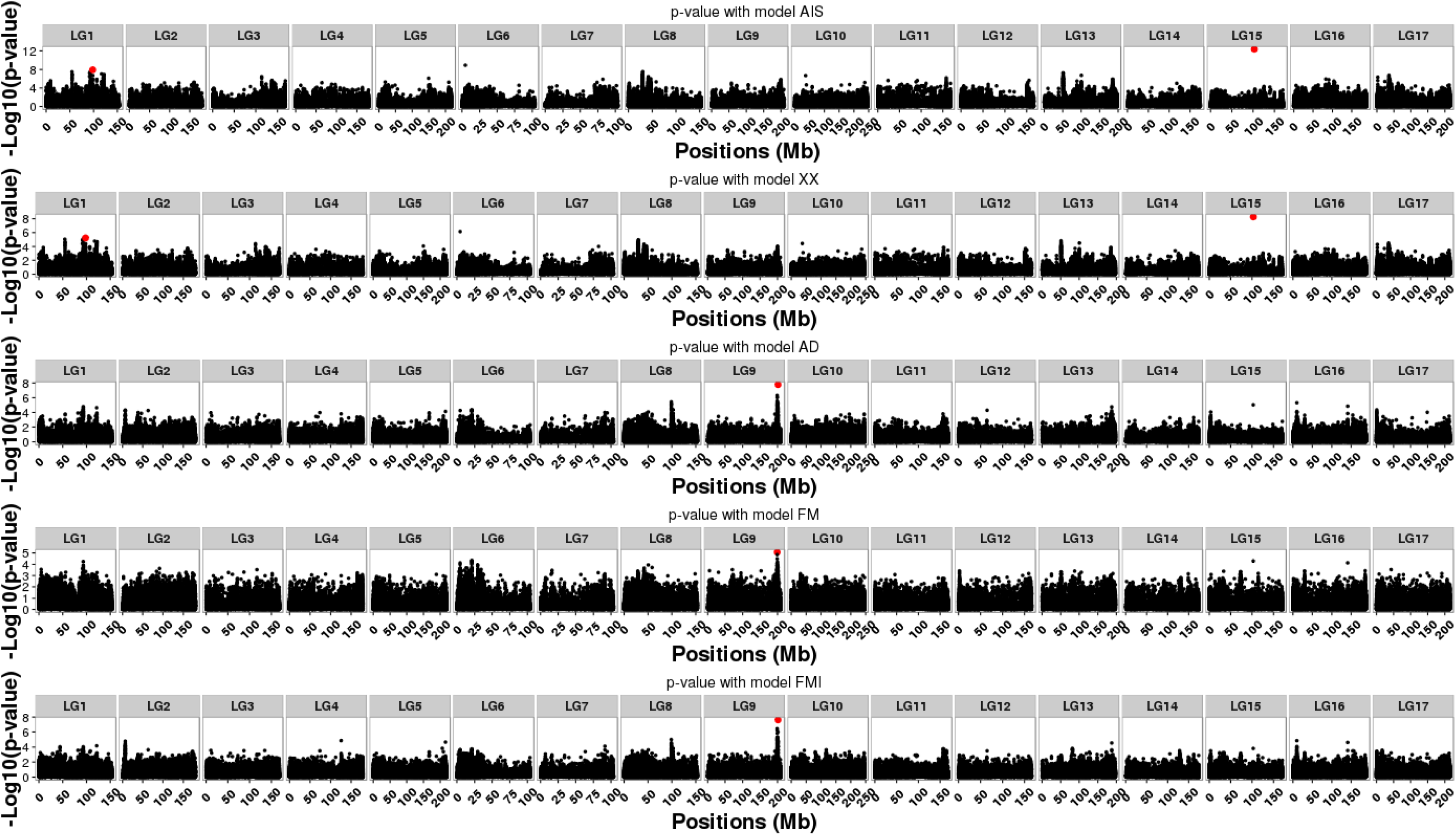
Manhattan plot for all models (*A*_*AIS*_, *A*_*XX*_, *AD*, *FM* and *FMI*) for the environment 13EX04. For each model, the p-values off all SNPs of the analysis are represented according to the position of the SNPs on the genome (in Mb). P-values are those of the step of the algorithm where all SNPs that are part of a QTL (red points) have been detected with the model of interest.

**Fig. S7:**
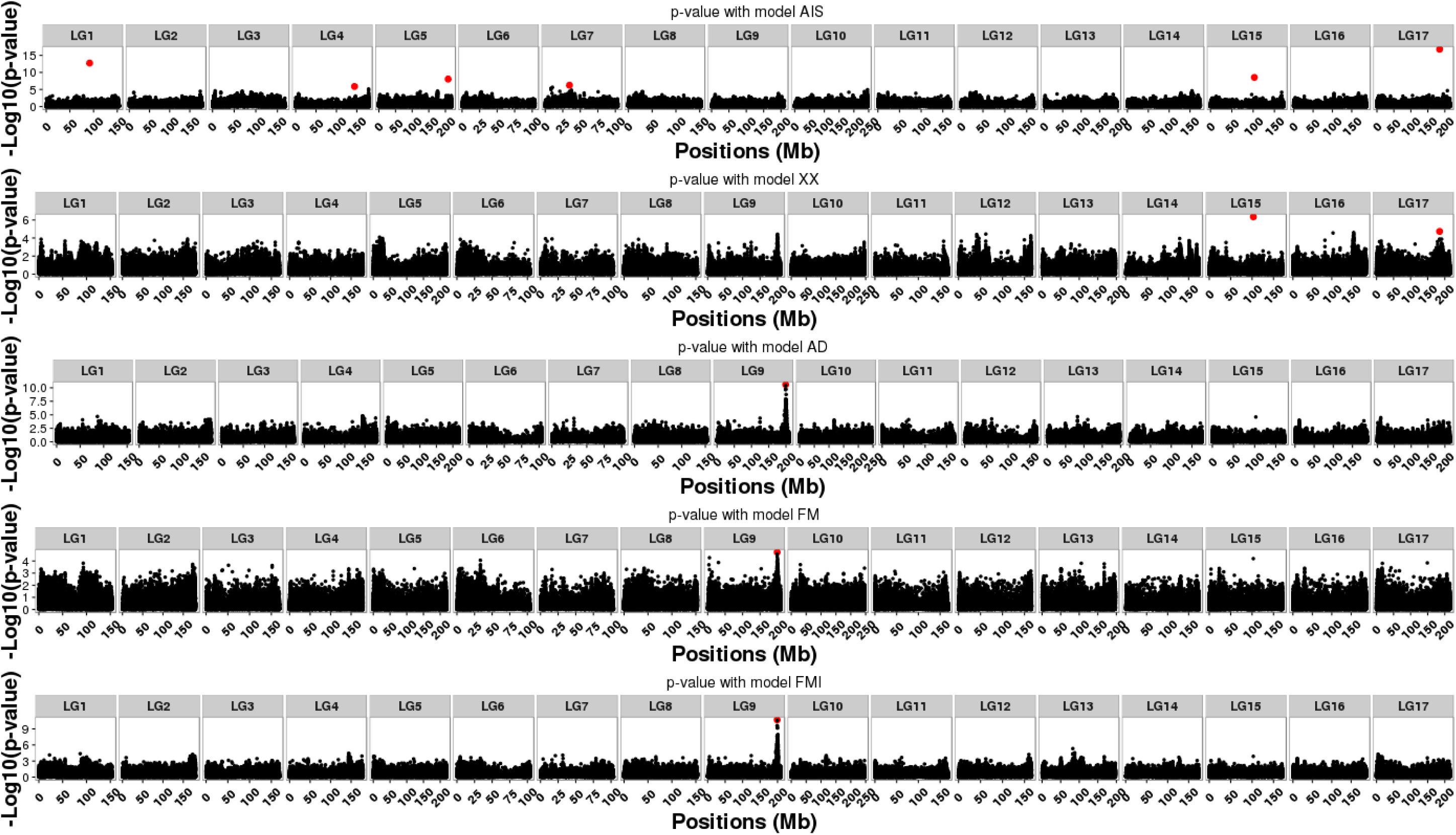
Manhattan plot for all models (*A*_*AIS*_, *A*_*XX*_, *AD*, *FM* and *FMI*) for the environment 13EX06. For each model, the p-values off all SNPs of the analysis are represented according to the position of the SNPs on the genome (in Mb). P-values are those of the step of the algorithm where all SNPs that are part of a QTL (red points) have been detected with the model of interest.

**Fig. S8:**
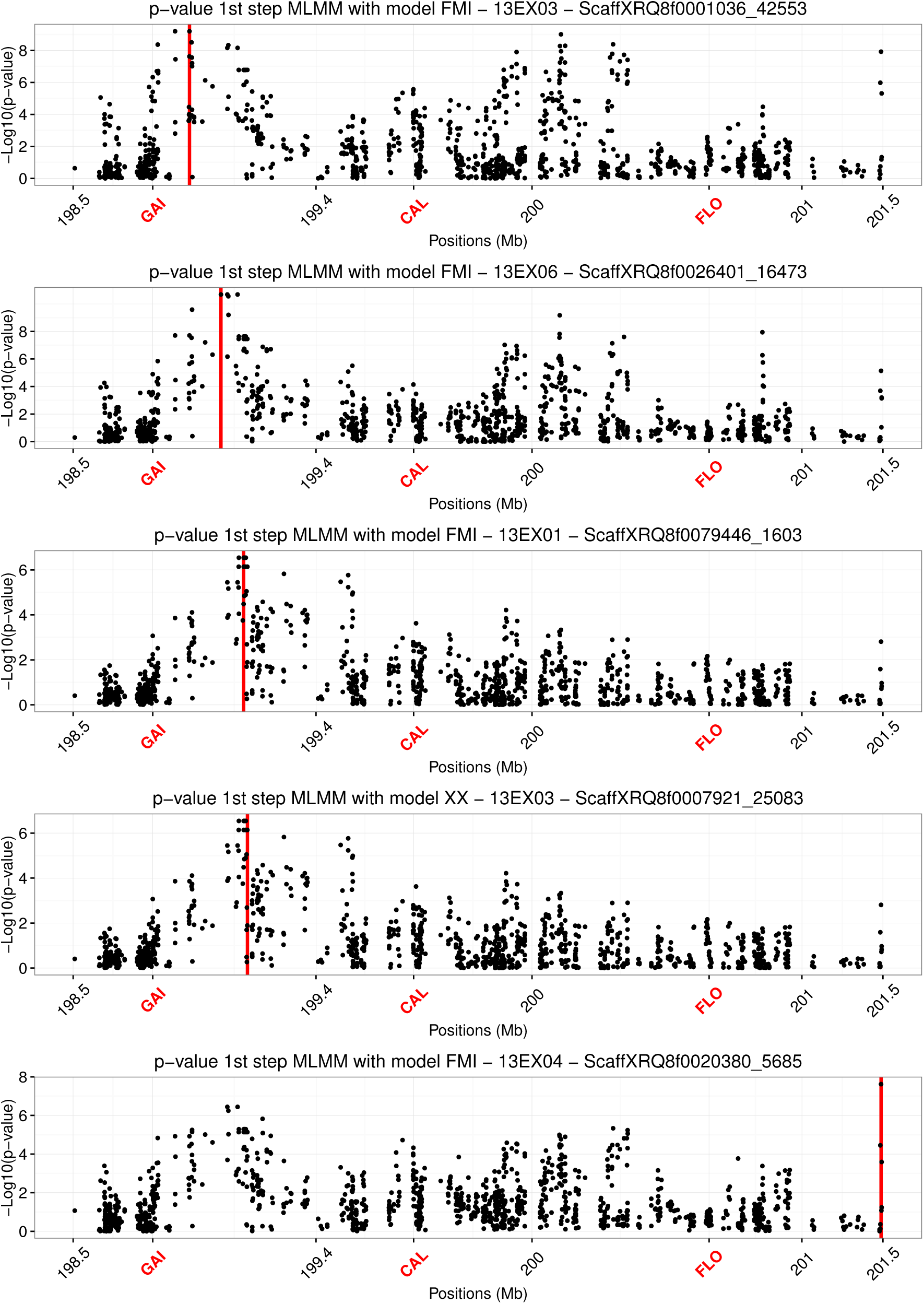
Manhattan plot of p-values of SNPs in QTL FT09.199. Combination of the association model and environment when significant association were found are illustrated. P-values are calculated in the corresponding model with the EMMAX model approximation. Red lines indicate the positions of associated SNPs. Gene involved in flowering process are positioned and indicated in red.

**Table S1:**
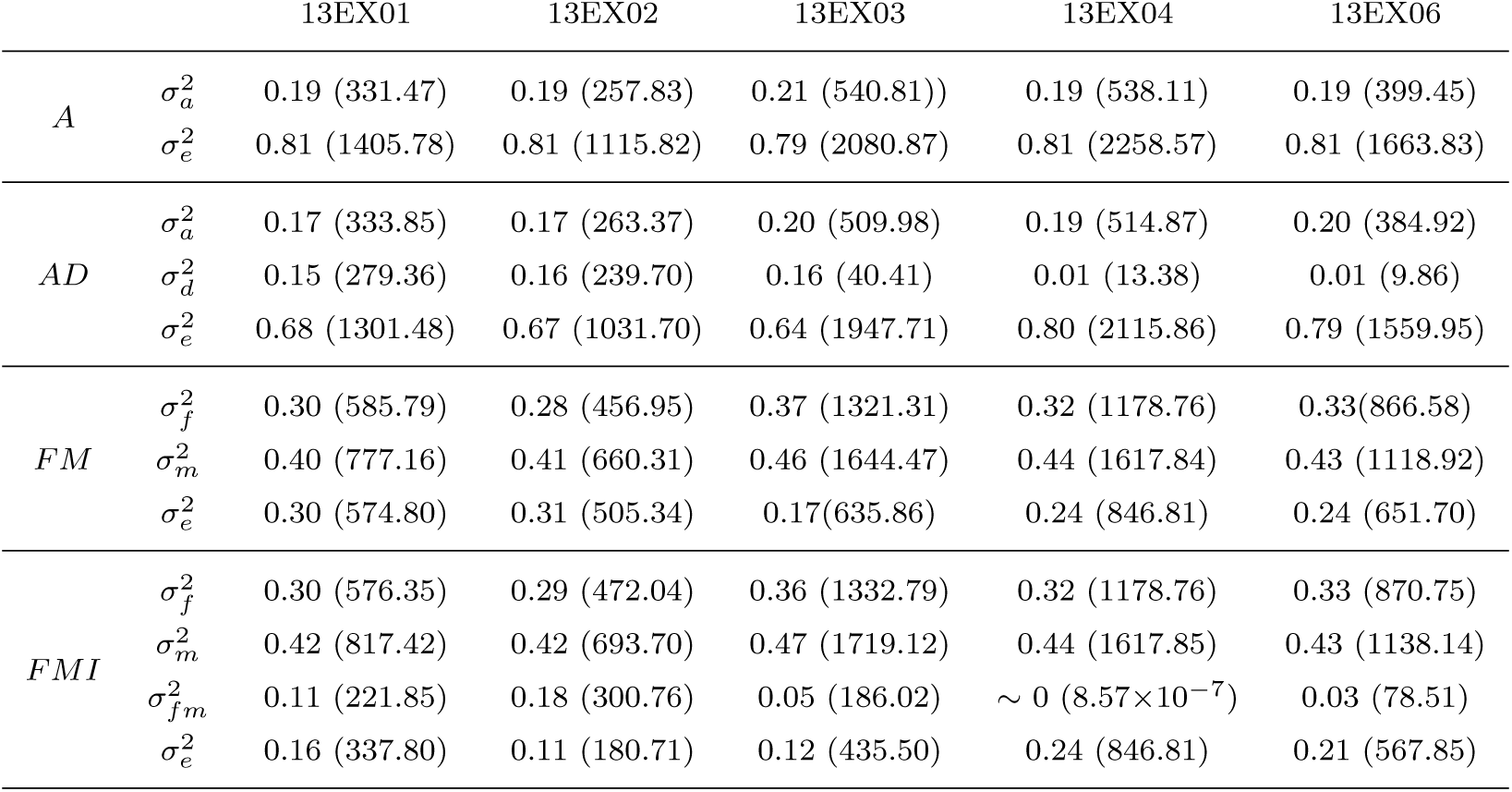
Parts of variance for the models *A* (*A*_*XX*′_), *AD*, *FM* and *FMI* for all the environments (13EX01 to 13EX06). 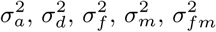 and 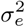 represent the additive, the dominant, the female, the male, the interaction between female and male, and the residual part of variance, respectively. The model used to compute theses parts of variance is the one without the effects of SNPs, using only kinships. The values of the variance components are shown in parentheses.

**Table S2:**
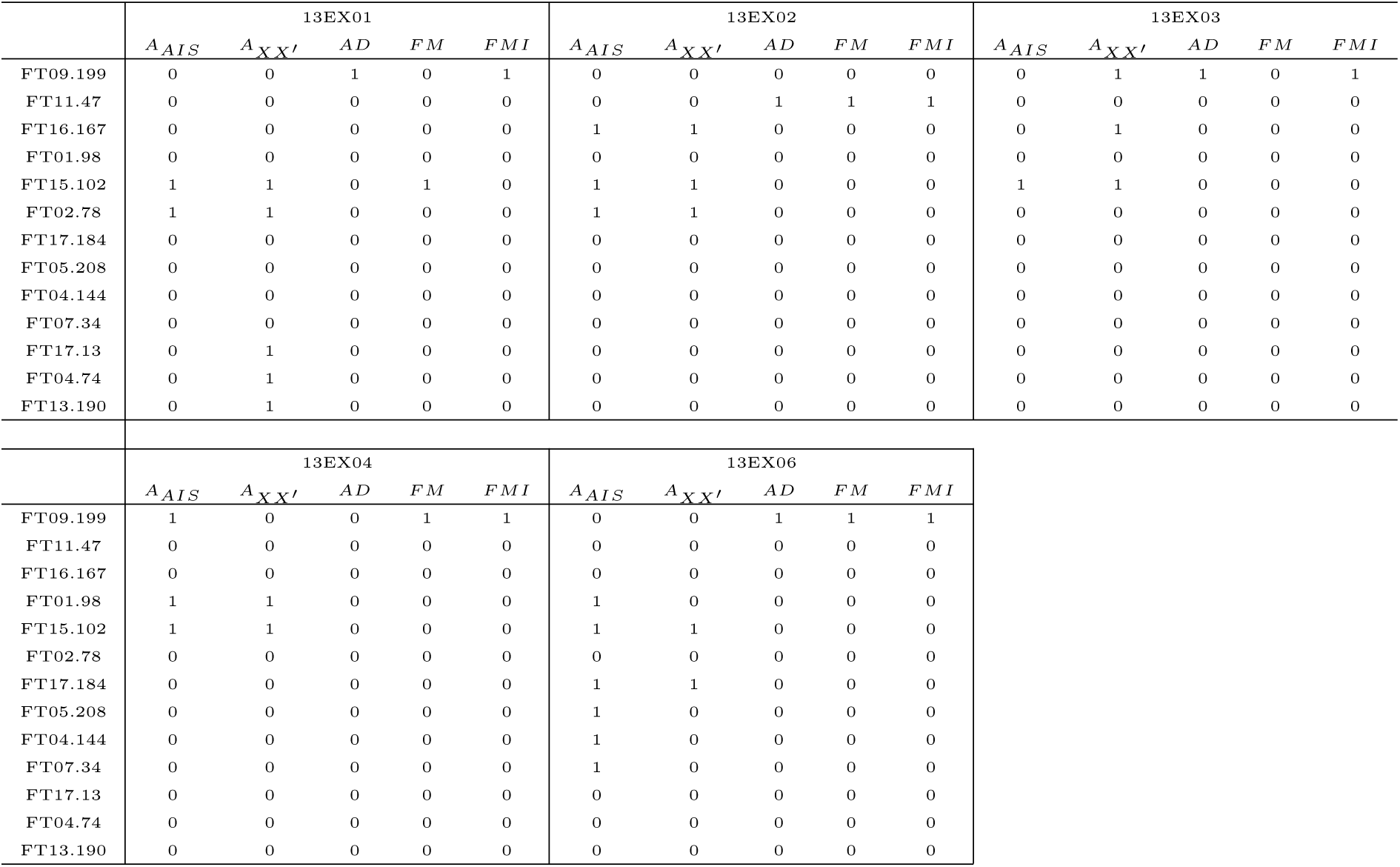
Summary of QTLs found for each model in each environment. “1” indicates that the QTL is associated to the flowering time in this environment with this model and “0” indicates no association.

